# Discovery of a cryptic pocket in the AI-predicted structure of PPM1D phosphatase explains the binding site and potency of its allosteric inhibitors

**DOI:** 10.1101/2023.03.22.533829

**Authors:** Artur Meller, Saulo de Oliveira, Aram Davtyan, Tigran Abramyan, Gregory R. Bowman, Henry van den Bedem

## Abstract

Virtual screening is a widely used tool for drug discovery, but its predictive power can vary dramatically depending on how much structural data is available. In the best case, crystal structures of a ligand-bound protein can help find more potent ligands. However, virtual screens tend to be less predictive when only ligand-free crystal structures are available, and even less predictive if a homology model or other predicted structure must be used. Here, we explore the possibility that this situation can be improved by better accounting for protein dynamics, as simulations started from a single structure have a reasonable chance of sampling nearby structures that are more compatible with ligand binding. As a specific example, we consider the cancer drug target PPM1D/Wip1 phosphatase, a protein that lacks crystal structures. High-throughput screens have led to the discovery of several allosteric inhibitors of PPM1D, but their binding mode remains unknown. To enable further drug discovery efforts, we assessed the predictive power of an AlphaFold-predicted structure of PPM1D and a Markov state model (MSM) built from molecular dynamics simulations initiated from that structure. Our simulations reveal a cryptic pocket at the interface between two important structural elements, the flap and hinge regions. Using deep learning to predict the pose quality of each docked compound for the active site and cryptic pocket suggests that the inhibitors strongly prefer binding to the cryptic pocket, consistent with their allosteric effect. The predicted affinities for the dynamically uncovered cryptic pocket also recapitulate the relative potencies of the compounds (τ_b_=0.70) better than the predicted affinities for the static AlphaFold-predicted structure (τ_b_=0.42). Taken together, these results suggest that targeting the cryptic pocket is a good strategy for drugging PPM1D and, more generally, that conformations selected from simulation can improve virtual screening when limited structural data is available.

## Introduction

Virtual screening is a common tool for identifying novel inhibitors of proteins with known structures.(Wallach et al., 2015; Lyu et al., 2019; Bender et al., 2021) Conventional, structure-based virtual high throughput screening approaches use an empirical-or force-field-based scoring function to dock ligands to mostly rigid receptors and rank compounds.(Trott and Olson, 2010) Docking to structures that deviate from the ligand-bound state can result in inaccurate predictions of the bound complex and poor compound ranking. For example, it is often difficult to recover active compounds when docking against ligand-free experimental structures (e.g., an *apo* state), or when the cognate ligand is small.(Abagyan et al., 2010) Even worse, experimentally derived structures are unavailable for many targets with disordered or flexible domains.

AlphaFold (AF) has the potential to accelerate drug discovery thanks to accurate structure prediction for such proteins.(Jumper et al., 2021) However, these are still just rigid structures, and their utility will be limited if they do not represent bound-like structures.(Vijayan et al., 2015; Wankowicz et al., 2022)

Phosphatases are a protein family with many potential therapeutic targets, but few are currently drugged (Mullard, 2018; Köhn, 2020) owing to a highly conserved and charged active site. Phosphatases are distinguished by different functional domains that can be exploited for the design of selective therapeutics (e.g., SH2 domain in SHP2(Chen et al., 2016)). Often, these domains are highly flexible.(Miller et al., 2022) Human protein phosphatase, Mg2+/Mn2+ dependent 1D PPM1D, also known as Wip1, is an important therapeutic target in oncology.(Pecháčková et al., 2017) PPM1D negatively regulates p53 and other components of the DNA damage response pathway.(Lu et al., 2008) Overactivation of PPM1D, either through duplication or loss of its degradation domain, is present in several human cancers, including breast cancer (Li et al., 2002), ovarian clear cell carcinoma (Tan et al., 2009), and brain cancers (Castellino et al., 2008).

Several allosteric inhibitors of PPM1D have been discovered through experimental screens(Gilmartin et al., 2014), but they remain difficult to improve upon because PPM1D has defied structure determination. A dual biophysical and biochemical screen targeting PPM1D revealed a novel class of inhibitors called the capped amino acids (CAA).(Gilmartin et al., 2014) These compounds selectively and non-competitively inhibit the phosphatase activity of PPM1D towards FDP and natural substrates. Efforts to crystallize PPM1D alone or PPM1D in complex with these inhibitors were repeatedly unsuccessful, likely due to a highly disordered loop or a flexible flap domain.

In the absence of this structural information, two distinct binding modes have been proposed based on indirect evidence. Photoaffinity labeling experiments suggested that the allosteric compounds bind at the PPM1D flap domain, in the vicinity of P219 and M236 (Fig. 1). (Gilmartin et al., 2014) In support of this model, the authors demonstrated that swapping the flap domain of PPM1D into another phosphatase rendered that protein sensitive to the PPM1D inhibitors. However, this finding was later disputed by several experiments that implicated the hinge domain in the binding of the allosteric compounds.(Miller et al., 2022) Deletion of the flap domain did not have an impact on the thermal shift, binding affinity, or the deuterium exchange profile caused by one of the allosteric compounds. Conversely, deletion of the hinge contributed to a substantial decrease in binding affinity and inhibition (i.e., an increase in IC50). Thus, the lack of experimental structures as well as competing binding modes makes PPM1D a uniquely challenging target for computational drug design.

**Figure 1:**
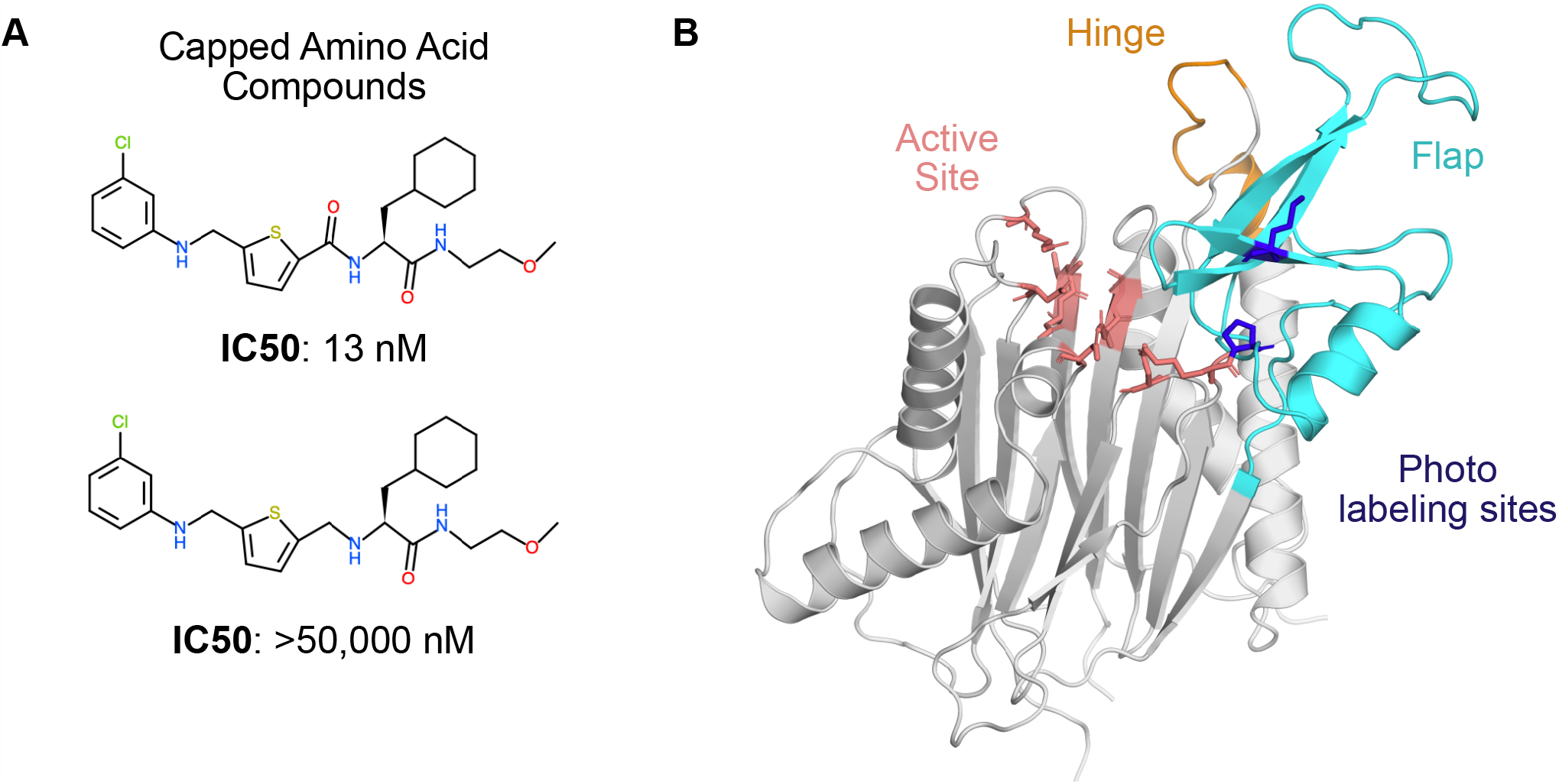
PPM1D phosphatase is allosterically inhibited by the capped amino acid (CAA) compounds, but the precise binding site is unknown. **A)** The capped amino acid compounds have a common amino acid-like substructure, and small differences in their chemical structure (i.e., the absence of a carbonyl) can contribute to very large differences in their potency. **B)** The AlphaFold-predicted structure of PPM1D highlights key regions that have been implicated in the binding of the capped amino acid compounds. The active site is shown in salmon sticks while two residues identified as proximal to the binding site based on photolabeling experiments are shown in blue sticks. The flap domain, a region hypothesized to be the primary CAA compound binding site, is shown in cyan. Another region hypothesized to be the primary CAA compound binding site, the hinge, is shown in orange.

Here, we use AlphaFold, molecular dynamics simulations (Karplus and McCammon, 2002; Hollingsworth and Dror, 2018), and machine learning to generate distinct conformations of PPM1D to investigate the molecular mechanisms of allosteric inhibition.

## Results

### PPM1D’s AlphaFold structure lacks high scoring pockets at the flap and the hinge

Given the lack of available PPM1D experimental structures, we first tested if a structure predicted by AlphaFold (AF) could help determine the preferred binding site for its allosteric inhibitors. The high accuracy of AF predictions(Jumper et al., 2021) suggests that structures predicted by AF can be used for determining binding sites and conducting virtual high throughput screening campaigns. Therefore, we analyzed the PPM1D AF structure to determine if there were binding sites with a high probability of ligand binding.

The PPM1D AlphaFold structure lacks clear pockets at the flap and the hinge, which are the two binding sites proposed in the literature. In contrast to previous homology models constructed for PPM1D, the AF structure of PPM1D includes a structured flap domain. The predicted local distance difference test (pLDDT) score, a useful proxy for how ordered a region is (Wilson et al., 2022), is high in the flap domain (Fig S1).

Despite the structured nature of the flap domain, there are few obvious pockets for an allosteric inhibitor to bind. Using the P2rank algorithm (Krivák and Hoksza, 2018), we evaluated pockets on the protein surface and found two pockets with high scores (Fig S2). One is at the active site, which cannot be the preferred binding mode for the capped amino acid compounds given the non-competitive nature of PPM1D inhibition. The second high scoring pocket is found opposite the flap domain where helix 323-326 and helix 347-360 interface with one of the β-strands in the PPM1D β-sandwich (Fig S1). This pocket has no overlap with either of the proposed binding sites found in the literature for the PPM1D allosteric compounds. Both the flap and the hinge lack high scoring pockets in their vicinity. Similarly, when we searched for pockets using the LIGSITE algorithm (Hendlich et al., 1997), we do not find pockets at either of the proposed binding sites (Fig S3). These findings suggest that the binding site of the allosteric inhibitors is possibly cryptic or transient, or simply not captured by the AlphaFold structure – thus posing a challenge for a successful docking campaign.

Hence, we decided to investigate whether molecular dynamics simulations might reveal cryptic pockets at the flap or the hinge.

### PPM1D *apo* simulations reveal a cryptic pocket at the flap-hinge interface

Next, inspired by recent success in capturing cryptic pocket formation in molecular dynamics simulations,(Hollingsworth et al., 2019; Sztain et al., 2021; Zimmerman et al., 2021; Cruz et al., 2022; Meller et al., 2022b, 2023) we tested whether simulations launched from the AF structure could reveal cryptic pockets that encompass the flap or the hinge. We used an adaptive sampling algorithm FAST (Zimmerman and Bowman, 2015) to search for cryptic pockets. FAST balances exploration with exploitation to efficiently search conformational space for conformations with desired traits. FAST does this by launching swarms of simulations and then selecting the most promising states as evaluated by an objective function for further simulations. In our case, we defined an objective function that included LIGSITE pocket volume to favor states with large pockets and another term to reward conformations which had been rarely observed (see Methods). Following each round of simulations, we created Markov State Models (MSMs) (Pande et al., 2010; Bowman et al., 2015) of the protein’s conformational ensemble after clustering conformations using C-α RMSD as a distance metric.

In our simulations, the flap domain is extremely dynamic, sampling closed and highly open conformations (Fig. 2A). An MSM-weighted distribution of flap domain to active site distances reveals two modes, one centered roughly on the distance found in the AF starting structure (∼23 Å) and another around 27 Å (Fig. 2A). In the closed conformations with a small active site-flap distance, the flap domain approaches a helix (residues 346-361) whose minimum distance to the flap domain in the AF structure is 11 Å (structure I in Fig. 2B, Fig. S4). This behavior is consistent with experiments which showed that flap deletion leads to an increase in deuterium incorporation, implying an increase in backbone solvent exposure, at peptides spanning residues 328-362.(Miller et al., 2022) Not only can the flap close in on the active site, it can also dissociate dramatically as seen in the long tail on the right of the active site-flap distance distribution (structure iii in Fig. 2B). In this extended conformation, K218 and other residues involved in substrate recognition are far from the active site (i.e., the distance between K218’s sidechain to D105’s sidechain grows from 9 Å in the AF structure to as much as 29 Å in simulations). The two peaks seen in the flap domain to active site distance distribution are consistent with both hydrogen deuterium exchange mass spectrometry and sedimentation velocity ultracentrifugation experiments(Miller et al., 2022), which showed that PPM1D exists in an equilibrium between two different flap domain conformations.

**Figure 2:**
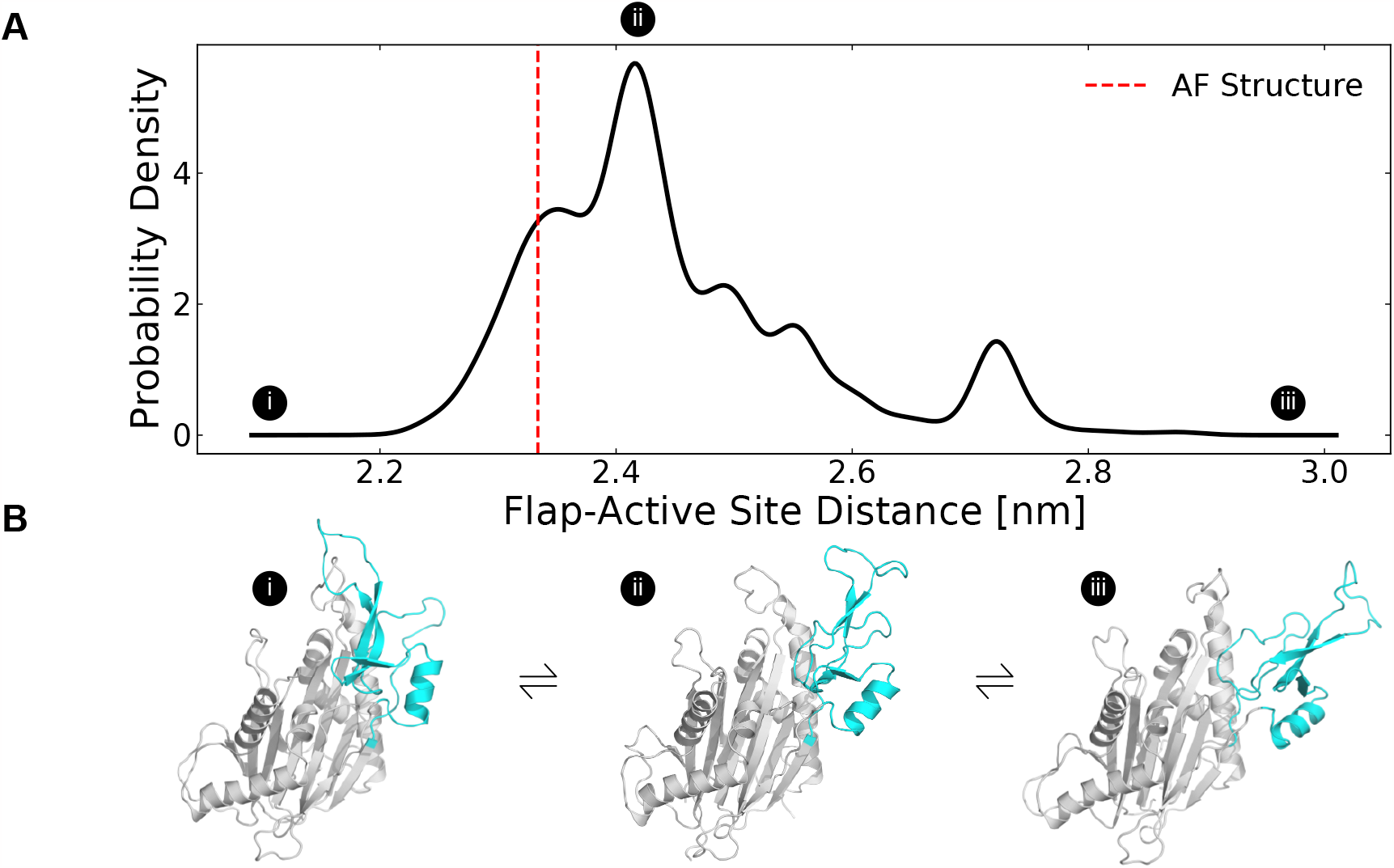
The distribution of flap domain to active site distances from MD simulations highlights that the flap is a highly flexible domain that can adopt more open conformations than seen in the AlphaFold-predicted structure. **A)** The MSM-weighted distribution of average distances between the flap domain (defined as residue 219-295) and the active site (residues 105, 192, 314, and 366) backbones shows two peaks as well as long tails that highlight low probability highly closed and highly open conformations. The dashed red line indicates the same distance measured for the AlphaFold-predicted structure. **B)** These structures depict a highly closed, an intermediate, and a highly open MSM cluster center. The flap domain is colored in cyan. Circles with Roman numerals indicate where these structures fall in the distribution.

The highly dynamic nature of the flap domain is not captured in the AlphaFold predictions. As predicted by the high pLDDT estimates for the flap domain, the β-strands in the flap remain structured as β-strands throughout the simulations (Fig. S5). However, neither AF’s pLDDT nor the predicted aligned error for the flap domain suggest that flap domain dissociation is possible or likely. We speculate that AF underestimates flap domain flexibility because it is trained with static structures from the Protein Databank (PDB), and thus simulations are a useful means to identify functionally important excited states.

Our simulations revealed a cryptic pocket at the flap-hinge interface between the two proposed binding sites. We calculated pockets for each structure in the MSM using P2Rank (see Methods). We then found the difference in each residue’s maximum ligand-binding probability in the ensemble and its ligand-binding probability in the AlphaFold structure. This analysis revealed that the flap domain, especially a flap domain loop (residues 276-290), is enriched for residues with large increases in ligand-binding probability (Fig S6, S7). To visualize this flap domain cryptic pocket, we found the simulation structure with the largest increase in predicted ligand-binding probability relative to the AlphaFold structure. This structure shows conformational changes in the orientation of the central β-strand in the flap as well as the loop spanning residues 269-295 (Fig 3A). Collectively, these lead to the formation of a deep pocket (Fig 3B, D) with a P2Rank-predicted ligand-binding probability of 0.87. There are other regions of the protein with increases in predicted ligand-binding probability, including the hinge (Fig S8) and the photoaffinity labeling sites, (Fig S9) but these increases are not as substantial as those in the flap domain loop. Taken together, these results suggested that relevant binding modes for the PPM1D allosteric compounds may be hidden in the ground state AlphaFold structure.

**Figure 3:**
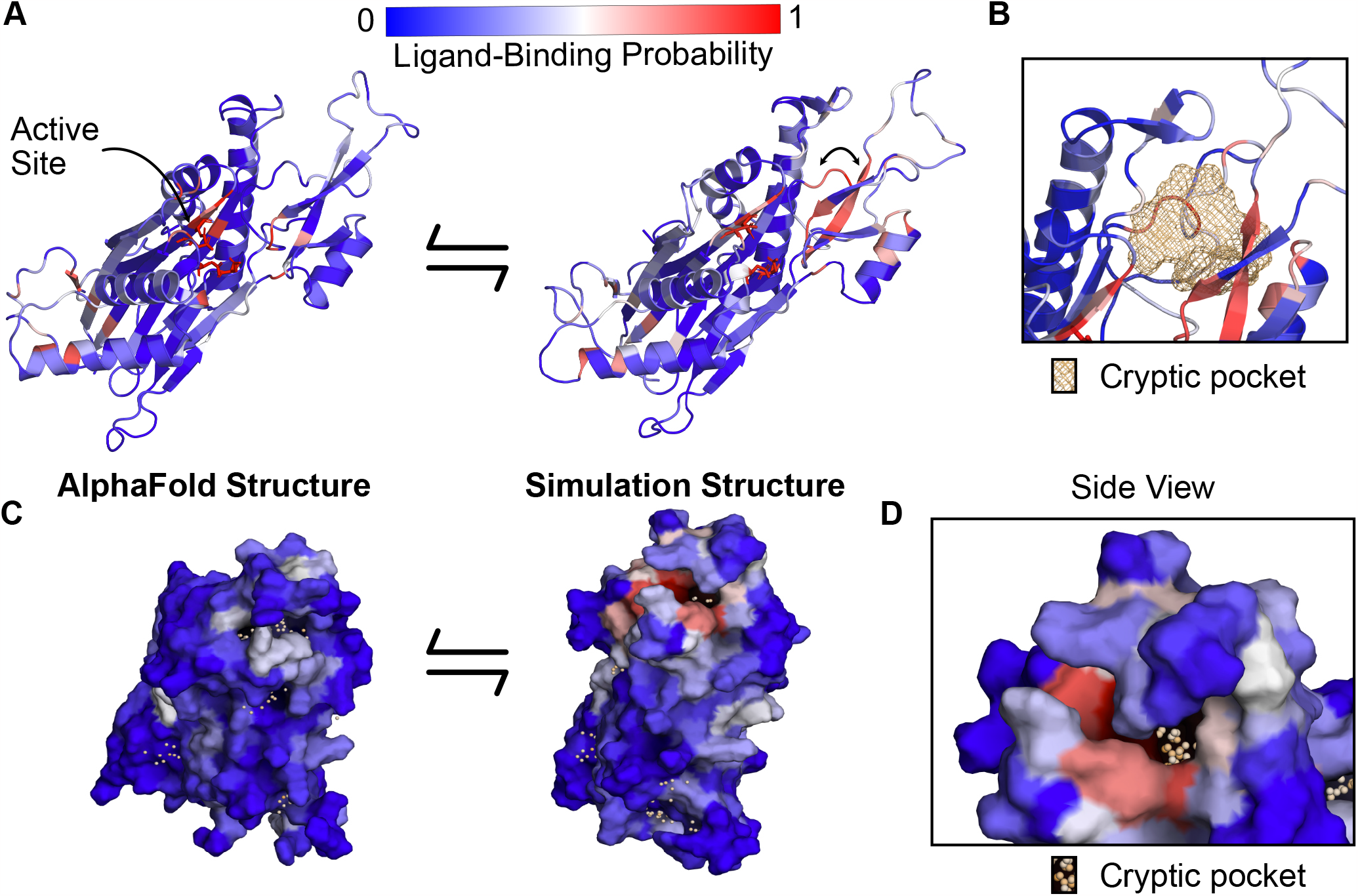
PPM1D *apo* simulations reveal a cryptic pocket at the flap-hinge interface. **A)** The AlphaFold-predicted PPM1D structure and a simulation structure where each residue is colored by its P2Rank-predicted ligand-binding probabilities show an increase in ligand-binding probability at the flap domain near the hinge. This simulation structure was selected because it had the largest increases in ligand-binding probability relative to the starting structure across the ensemble of states. Active site residues are shown in sticks. Arrow indicates the backbone motion that is required to form the cryptic pocket. **B)** Mesh representation of the cryptic pocket shows that it forms between a flap domain loop (residues 276-279), two of the β-strands in the flap (residues 243-247 and 268-271), and a flap domain helix (residues 227-234). **C)** Surface representation looking onto the AlphaFold structure and the open simulation structure highlights that a deep trench forms between the flap domain and hinge.The surface is colored by P2Rank-predicted ligand-binding probability. **D)** A zoom-in of the surface representation of the open state reveals that the cryptic pocket lies in a deep groove. The orange spheres are the pocket grid points identified by P2Rank.

### The AtomNet PoseRanker neural network predicts a single preferred cryptic binding site between the flap and hinge

To help determine which cryptic site was the most likely binding site, we docked the PPM1D allosteric compounds across the ensemble of structures in our MSMs.

Traditional rigid body docking can often produce high quality poses (root mean square deviation from a crystal pose less than 2 Å), but these methods struggle to rank the poses correctly (Su et al., 2019); the highest quality poses rarely correspond to the highest scoring poses. To circumvent this limitation, deep learning methods often re-rank conventional docking poses and achieve improved performance. We used one of these methods, AtomNet PoseRanker (ANPR), to re-rank the poses from molecular docking. (Stafford et al., 2022) ANPR was trained on existing data on the PDB and demonstrated to have an implicit understanding of physical interactions and protein dynamics. ANPR is trained as a binary classifier, and outputs a probability score between 0 and 1 (scores greater than 0.5 are usually indicative that ANPR has confidence that the pose in question is of high quality). We hypothesized that correctly assigned binding sites for ligands would admit better poses than incorrect sites. We therefore used ANPR scores to evaluate and identify the most likely binding site of the PPM1D allosteric inhibitors. We expected the most likely binding site to have higher ANPR scores across the simulated conformations with a relevant cryptic pocket.

We docked compounds to all states from the PPM1D MSMs using CUina (Gniewek et al., n.d.; Stafford et al., 2022), a GPU-efficient implementation of smina (Koes et al., 2013), and evaluated the quality of the resulting docked poses with ANPR. For every state from the MSM, we used P2Rank to identify possible binding sites in that state’s representative structure. A significant number of conformations presented a cryptic pocket between the hinge and the flap. A smaller number of conformations presented a pocket almost exclusively at the hinge. We used the pockets identified by P2Rank to design a box centered around these pockets. We padded the box by 5 Å on each dimension, and we used that box to define the search space of our molecular docking runs. As a control, two additional bounding boxes were created for the active site and photolabeling site described in the Gilmartin publication by defining the boundaries based on the catalytic residues or the photo labeling residues respectively. These boxes were also padded by 5 Å in each dimension (see Methods). In total, we docked nine capped amino acid compounds against four possible sites (two proposed sites around the hinge, the photolabeling site, and the active site as a negative control). These compounds were docked against all MSM states where the relevant cryptic pocket was detected by P2Rank. For each compound + binding site pair, we re-ranked the top 64 poses (as ranked by the vina scoring function) using ANPR. The pose for each compound and binding site with the highest ANPR ranking was selected for subsequent analyses. Interestingly, none of the poses where PPM1D allosteric compounds were docked to the AF structure scored above 0.5, indicating that these were unfavorable poses (Table S1). This corroborates our pocket assessment results, suggesting that the static AF structure is not amenable to docking of the PPM1D allosteric inhibitors.

Across the PPM1D MSM ensemble, we found that ANPR assigns the highest scores to poses where the compounds bind between the flap and hinge. For each compound, we assessed which poses were given a ANPR probability score greater than 0.5. We defined those as predicted high-quality poses. We found that residues found at the interface of the hinge and flap domain are most likely to make contacts with high-quality poses (Fig. 4A). Specifically, residues in the flap domain loop from D277 to V289 are most likely to form contacts with these poses. When we overlayed all high-quality poses of the compounds onto the AF starting structure, we found that they cluster in a single region between the flap and hinge (Fig S10). Next, we classified poses by the protein contacts that they form into the following categories: flap domain only, hinge only, flap-domain interface, and active site (see “Pose classification” in Methods). There are no high-quality poses that form contacts only with the hinge and rarely did any high-quality poses form contacts with the active site. This is true across all compounds. Considering that the PPM1D allosteric inhibitors are non-competitive, our negative control results (docking against the active site) bolster our confidence that the ANPR probability scores can distinguish between correct and incorrect sites. We used the equilibrium probabilities from the MSM to calculate a weighted average of the ANPR score across the PPM1D ensemble (Fig. S12). We find that the ensemble-weighted ANPR probability is highest at the flap domain and flap-hinge interface (Fig. 4B, S13). Thus, these ANPR predictions strongly suggest that PPM1D allosteric compounds bind between the flap and hinge.

**Figure 4:**
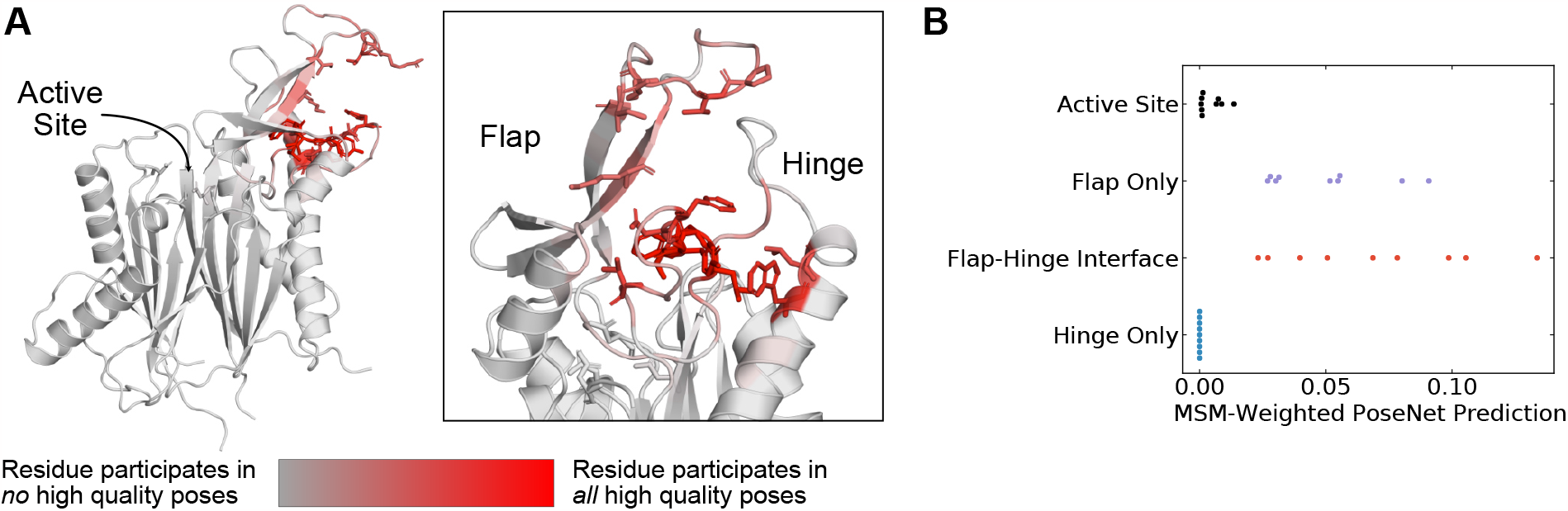
The AtomNet PoseRanker neural network predicts that poses found at the flap-hinge interface are more crystal-like. **A)** A PPM1D AlphaFold structure colored by the frequency with which residues participate in high-quality poses indicates that residues needed for high-quality poses are found at the flap-hinge interface. Residues in dark red most frequently contact the GSK2830371 compound in its high-quality poses. High-quality poses were those poses that received a PoseRanker score of 0.5 or higher. A contact was defined when a ligand heavy atom was within 4 Å of a protein heavy atom. **B)** The MSM-weighted AtomNet PoseRanker (ANPR) predictions across different binding sites show that the flap-hinge interface receives higher ANPR scores. Each point represents a different CAA compound. When there were multiple poses in one of the binding site categories, we selected the pose with the highest ANPR score. We defined the hinge as residues 150-166 and the flap as residues 219-295.

### Combining MSM-docking with pKi predictions from a neural network accurately ranks compounds

While an estimate of pose quality might be helpful in virtual screening, the decision to select compounds for synthesis and testing with *in vitro* assays relies on an estimate of a compound’s bioactivity or affinity. The deep learning-based pKi predictor AtomNet has been shown to be physics-aware and to be sensitive to pose perturbations.(Gniewek et al., n.d.; Wallach et al., 2015) Considering that the CAA compounds have known affinities, we can assess whether MSM-docking (Meller et al., 2023) can have an impact on the retrospective performance of the AtomNet pKi predictor.

We applied the AtomNet pKi predictor to each of the docked poses in our MSM ensemble. The AtomNet pKi predictor was trained using a combination of public and proprietary structural data. It outputs a value for the predicted pKi of a compound for a particular target given a particular pose provided as input. We docked each compound to several sites for each structure in the ensemble. We used the ANPR score to select the highest scoring pose per compound-structure pair in the ensemble (Fig. 5A). We then passed that compound-state pair as input to the AtomNet pKi predictor, resulting in one prediction of the compound’s potency per MSM state.

**Figure 5:**
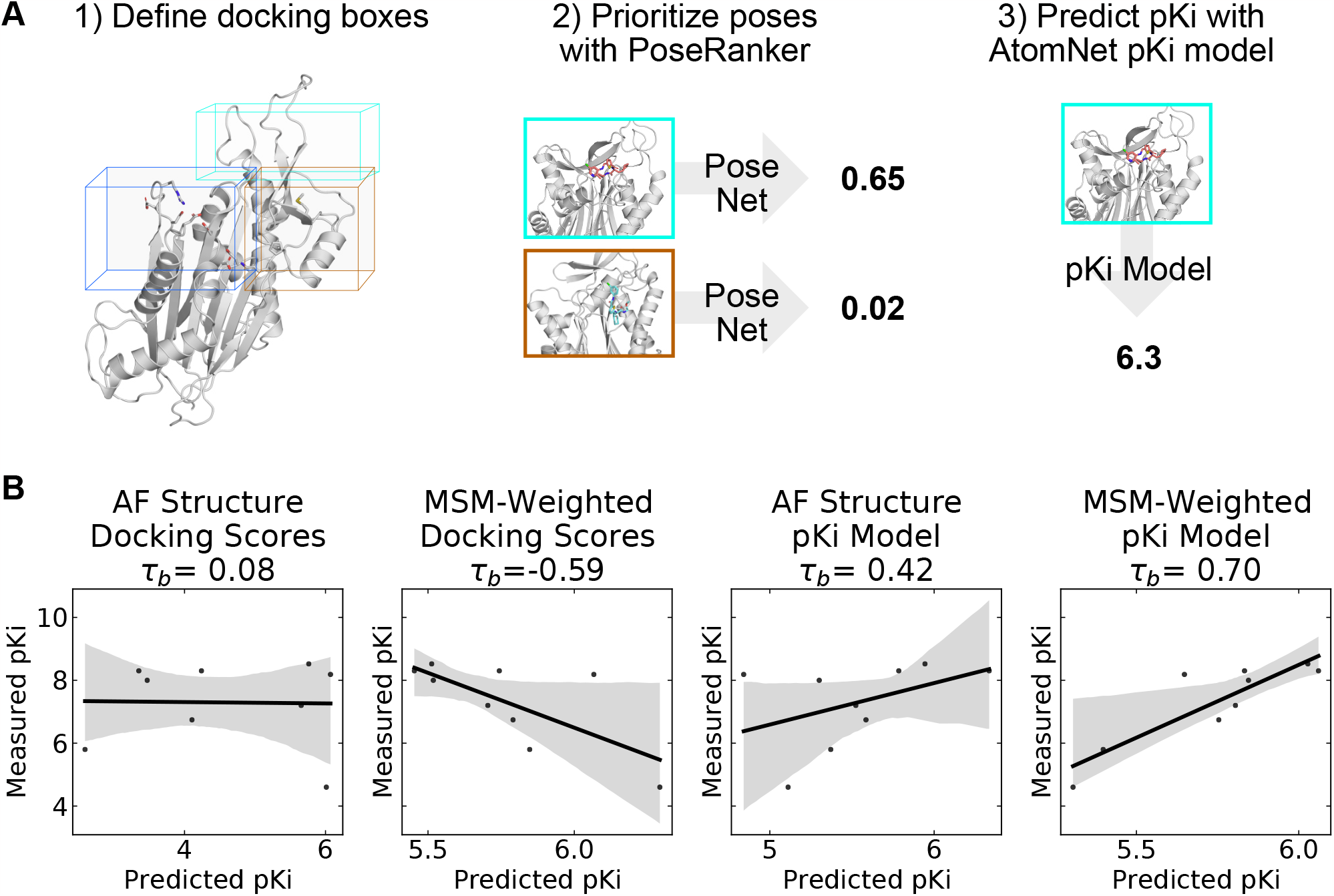
A neural network trained to predict pKi accurately ranks allosteric compounds by potency when applied to structures from a PPM1D ensemble. **A)** Schematic highlighting the procedure that was used for selecting a single pose for each PPM1D cluster center in the MSM. For each MSM cluster center, we defined multiple docking boxes based on the active site, residues involved in photolabeling experiments, and P2Rank pockets at the flap and hinge. After performing docking, we selected a best pose per MSM state using the PoseRanker neural network. Finally, we fed this best docked pose to the AtomNet pKi predictor. **B)** MSM-weighting of the pKi predictions from the AtomNet pKi predictor outperforms docking-based methods as well as a single pKi prediction based on the AlphaFold-predicted structure. For each scatter plot, we show the line of best fit in black as well as the 95% confidence interval based on bootstrapping in translucent grey bands. We report the Kendall rank correlation coefficient, a statistic that measures the ordinal association between the predited pKi and the measured pKi and whose maximum value is 1.

We find that taking an ensemble perspective that accounts for cryptic pockets outperforms results for the static AF structure. We first established a baseline by evaluating how well docking scores rank PPM1D allosteric compounds by potency. Docking scores for the AF structure alone and MSM-weighted docking scores for the ensemble (see Methods) generated very poor predictions of compound potency, demonstrating that ranking these compounds is a non-trivial task. In fact, compounds with better docking scores were less potent in general (Kendall τ_b_=-0.59, Fig 5B); we noticed negative correlation between docking scores and their measured potency. On the other hand, the AtomNet pKi predictor ranks more potent compounds higher using docked poses against the AF structure alone (τ_b_=0.42, Fig. 5B). The ability to rank compounds based on their predicted affinity further improves when we dock to all MSM states and weight the pKi predictions based on the equilibrium probability of each state (see Methods). Indeed, we achieve an impressive τ_b_ of 0.70 when using MSM-weighted pKi predictions (Fig. 5B). Thus, combining MSMs with the AtomNet pKi predictor may improve the performance of virtual screening.

## Discussion

Protein phosphatases are a challenging class of drug targets that broadly illustrate the advantages of using allosteric compounds.(Köhn, 2020) There are nearly 200 phosphatases in the human genome, and many are implicated in human diseases, including diabetes (Krishnan et al., 2018), neurodegeneration (Vieira et al., 2017), and multiple cancers (Pecháčková et al., 2017). Phosphatases are downstream targets of several signaling pathways that integrate various cellular signals.(Lu et al., 2008) This suggests that targeting of phosphatases may be useful across numerous cancer subtypes caused by mutations of upstream proteins or in cases where tumors develop resistance to upstream therapies. However, to the best of our knowledge, there are no approved therapies that target phosphatases. Previous drug discovery efforts have focused on active site inhibitors. Targeting the active site has proved challenging because high sequence conservation limits the selectivity of compounds. Furthermore, compounds targeting the active site need to be highly charged, limiting their bioavailability. Hence, allosteric compounds, like the CAA compounds that target PPM1D and novel allosteric inhibitors of SHP2 (Chen et al., 2016), may be needed to successfully inhibit phosphatases in clinical settings.

Definitively establishing the binding site of the PPM1D allosteric compounds remains challenging, but our results predict a plausible binding site that agrees with most previous experiments. Photoaffinity labeling experiments and flap swap experiments, which showed that introducing the PPM1D flap domain can sensitize other phosphatases to the PPM1D allosteric inhibitors, strongly implicate the flap domain as the primary compound binding site. Our proposed binding site at the flap-hinge interface is consistent with these results. Though our proposed binding mode does not directly involve the points of covalent attachment (i.e., P219 and M236), we speculate that the large photoactivatable benzophenone groups that were added to the compound scaffold enable compounds with these groups to bind at our proposed site but still reach these residues. Furthermore, Gilmartin et. al. showed that residues 247-268 in the flap are not essential for PPM1D allosteric compound binding.(Gilmartin et al., 2014) Consistent with these results, our proposed binding site does not involve these residues with the minor exception of K247. On the other hand, Miller et. al. demonstrated that deletion of the hinge causes a ∼1000-fold decrease in binding affinity and a 100-fold increase in IC50 for one of the allosteric compounds. Our proposed binding site has substantial involvement from hinge residue L157 and an adjacent residue W154. As a result, our proposed binding site is consistent with the hinge deletion experiments. However, given that residues in the flap, especially residues D277 to V289, are commonly involved in high-quality poses, we cannot explain why Miller et. al. report that flap deletion (specifically residues 219-287) has no effect on binding affinity or binding kinetics. We speculate that it may be possible for the allosteric inhibitors to bind even when most of the flap is deleted, but our analysis suggests further experiments are needed to disentangle the relative contributions of the flap and hinge to compound binding.

Furthermore, our results highlight the advantages of explicitly accounting for protein conformational heterogeneity when using deep learning methods for predicting compound affinity. The AtomNet pKi predictor is designed and trained to be pose-sensitive.(Gniewek et al., n.d.; Wallach et al., 2015; Stafford et al., 2022) Its performance at ranking compounds varies widely between target structures in the MSM (Fig S14). We noticed that even when the poses are likely of poor quality (e.g., the AF structure where the cryptic pocket is not present), we still often see relatively good predictive performance for the pKis. While some of the predictive power of the AtomNet pKi predictor is driven by the pose, we hypothesize that the ligand features might also play a part in and influence the predicted pKis that AtomNet pKi predictor outputs. For the cases where the pose is poor (e.g., docking against AF structure), we get a baseline for how well a ligand-based model would perform. The boost in performance seen with MSM-docking is likely due to better poses resulting from docking to structures with open cryptic pockets.

Our results also show that MSMs can address some of the limitations of rigid docking against AlphaFold predicted protein structures. Rigid docking has lower performance when the protein structure(s) being used for docking corresponds to an *apo* or unbound state.(Abagyan et al., 2010) Deep learning-based (DL-based) protein structure prediction methods like AlphaFold, are trained using all available data on the PDB, and there is data to support that output structures are somewhere in between *apo* and *holo*.(Saldanõ et al., 2022) Docking efforts against AlphaFold structures show lower performance than against *holo* structures available on the PDB.(Díaz-Rovira et al., 2022; Wong et al., 2022) Here, we show that this can be mitigated by considering conformational heterogeneity using MSMs. Using a highly flexible system, we can sample conformations and identify cryptic pockets that can be successfully used in downstream virtual screening applications. While our work was based off a single AF structure as a starting point, we are aware of efforts to use these DL protein structure prediction tools to sample multiple conformations, thus better capturing protein flexibility.(Meller et al., 2022a; Saldanõ et al., 2022) To our knowledge, these methods have not been compared against MSM approaches and more research would be needed before conducting a similar analysis as described herein with a DL-generated structural ensemble.

Despite these encouraging results, there are notable limitations to our approach.

Firstly, most of our pKi analyses included nine capped amino acid compounds. This is not a particularly large dataset, and we acknowledge that this is somewhat restrictive in terms of establishing robust statistical significance for our results. Ranking based on docking scores output by CUina does suggest that this is not a trivial ranking problem, and that achieving good predictive performance at random, despite the small data set size, is statistically unlikely. While in an ideal scenario we would hope to have a larger number of data points to validate our findings, affinity data is often relatively sparse at early stages of the pharmaceutical pipeline, so estimating the performance of virtual screening can be difficult. Secondly, our data suggests that the AtomNet pKi predictor tends to regress to the mean. Even though the ranking metrics are good, the dynamic range of predicted vs. observed pKis differ significantly. We hypothesize that this is likely due to a data imbalance in the training data of the AtomNet pKi predictor, as data points in the extremes of the pKi distribution (either very high or very low) are rare, and our sampling strategy during training does not stratify on that property. Still, given that model accurately ranks compounds by potency, our approach represents a promising strategy for novel virtual screening campaigns.

## Conclusions

In summary, we have uncovered a cryptic pocket at the PPM1D flap-hinge interface that improves the ability to predict the potency of PPM1D inhibitors. AlphaFold predicts a PPM1D structure that lacks high scoring allosteric pockets at proposed binding sites based on an analysis conducted using the P2Rank and LIGSITE pocket detection algorithms. Though the AF-predicted structure lacks allosteric pockets, molecular dynamics simulations of ligand-free PPM1D capture a cryptic pocket at the flap-hinge interface. A neural network trained to evaluate the quality of docked poses predicts that this site is the most likely binding mode for the PPM1D allosteric inhibitors. Finally, by docking compounds to this pocket and using a structure-based pKi predictor, we demonstrate that aggregating pKi predictions across a MSM is superior at ranking compounds than using docking scores or using the single predicted AlphaFold structure. Thus, our methodology provides a promising template for structure-based drug discovery and *in silico* binding site prediction.

## Methods

### Molecular Dynamics Simulations

The AlphaFold predicted structure (AF-O15297) was used as an initial structure for PPM1D simulations since no structures were available in the PDB. However, because several PPM1D domains (C-terminus domain and an internal loop stretching from residue 39 to 92) are predicted to be disordered (pLDDT < 70) and because we were primarily interested in flap domain dynamics, we removed residues 39-92 and truncated the C-terminus (residue 396-end).

GROMACS (Abraham et al., 2015) was used to prepare and to simulate PPM1D using the CHARMM36m force fields(Huang et al., 2016). The protein structure was solvated in a dodecahedral box of TIP3P water (Jorgensen et al., 1983) that extended 1 nm beyond the protein in every dimension. Thereafter, sodium and chloride ions were added to the system to maintain charge neutrality and 0.1 M NaCl concentration. The system was minimized using steepest descents until the maximum force on any atom decreased below 1000 kJ/(mol x nm). The system was then equilibrated with all atoms restrained in place at 310°K maintained by the Bussi-Parinello thermostat (Bussi et al., 2007) and the Parrinello-Rahman barostat (Parrinello and Rahman, 1998).

Production simulations were performed in the CHARMM36m forcefield. Simulations were run in the NPT ensemble at 310°K using the leapfrog integrator, Bussi-Parinello thermostat, and the Parrinello-Rahman barostat. A 12 Å cutoff distance was utilized with a force-based switching function starting at 10 Å. Periodic boundary conditions and the PME method were utilized to calculate the long-range electrostatic interactions with a grid density greater than 1.2 Å^-3^. Hydrogen bonds were constrained with the LINCS algorithm (Hess et al., 1997) to enable the use of a constant integration timestep of 2 fs.

### Adaptive Sampling

We used the Fluctuation Amplification of Specific Traits (FAST) algorithm (Zimmerman and Bowman, 2015) to explore a diverse ensemble of states with cryptic pockets. We used an objective function that rewarded states based on their total pocket volume as measured by LIGSITE.(Hendlich et al., 1997) The following LIGSITE parameters were used: a minimum rank of 7, a minimum cluster size of 3, and a probe radius of 0.14 nm. Our ranking function also included a term that penalizes states conformationally similar to others already selected (the width parameter for this term was 1.5 times the cluster radius)(Zimmerman et al., 2017). K-centers clustering was performed after each round with the RMSD of C-alpha positions of the entire protein as the distance metric. We set a cluster radius of 0.2 nm RMSD as a cutoff.

### P2Rank Pocket Detection

We used P2Rank v2.4 (Krivák and Hoksza, 2018) with default parameters to identify pockets across all of the representative states (cluster centroids) from our simulations.

For subsequent analyses, we consider only pockets with a permissive pocket probability (as output by P2Rank) greater than 0.2.

### Docking

We docked compounds using a proprietary GPU-enabled docking engine, CUina. CUina(Stafford et al., 2022) is a proprietary implementation of smina (Koes et al., 2013), which has been parallelized and refactored to operate more efficiently on a GPU. The scoring function (Vina scoring function) and sampling routines of CUina are analogous to those in smina.

CUina requires a bounding box to restrict its search space. We defined four bounding boxes representing each of the three proposes binding sites for CAA compounds, and one negative control (active site). For the first two boxes, we used the coordinates of the pockets identified by P2Rank in the vicinity of the flap or the hinge of PPM1D (where available). The minimum and maximum coordinates of the voxels output by P2Rank were used to define the box, and we padded these coordinates by 5 Å along each dimension. A third box was defined using the coordinates of the two residues (P219 and M236) that were part of the photolabeling experiment described by Gilmartin et al. The fourth and final boxed was defined based on the active site: we used the coordinates of all the catalytic residues to define the box. The box boundaries were calculated by taking the minimum and maximum coordinates of all photolabeling or catalytic residues and padding by 5 Å along each dimension.

We docked nine CAA compounds to all states (i.e., a representative structure for each MSM state) resulting from the MSM effort described above. For each compound, we dock the best (minimized) ligand conformation against all four proposed binding sites. In the MSM states where P2Rank failed to identify one of the pockets, docking against that pocket was omitted.

For each docking operation corresponding to a binding site + MSM representative structure + compound, we output 64 poses and imposed a 1 Å RMSD similarity cutoff, thus ensuring that the poses output are sufficiently different from one another.

#### Pose classification

Following docking, poses were classified based on the contacts that they formed. Specifically, we found residues whose heavy atoms were within 4 Å of a ligand heavy atom. Next, we classified poses into the following categories based on their list of contact residues: flap domain only, hinge only, flap-domain interface, and active site. The active site was defined as residues 18, 22, 23, 105, 106, 192, 218, 314, and 366 based on the annotation in (Gilmartin et al., 2014); the flap domain was defined as residues 219-288; and the hinge domain was defined as residues 150-167, which includes both a loop and half the helix spanning residues 136-158. If the compound made contacts with both a hinge domain and a flap domain residue, it was classified as binding in the flap-hinge interface.

### pKi Model Predictions

We used AtomNet’s pKi predictor to perform pKi predictions using the poses generated and selected by our pose generation pipeline (CUina + ANPR). AtomNet’s global pKi model uses a graph-based convolutional neural network to regress over pKi.

#### Data

This model was trained using a combination of public and proprietary data, spanning more than 4,000 targets for which activity measurements were available. In total, several million activity data points were used to train the model. PPM1D was not part of the training data for the model, but the training set did include a number of other phosphatases.

#### Architecture

AtomNet’s global pKi model uses the GRAPHite architecture (previously described in (Stafford et al., 2022). The GRAPHite architecture is a directed Graph Convolutional Network (GCN) comprised of four graph convolutional layers. The first two layers include both ligand and receptor features, whereas the last two layers are ligand-only. Nodes in the graph represent ligand and receptor atoms. Only receptor atoms within 7 Å of any ligand atom were used as part of the graph. Edges were defined by atoms within 4 Å of each other and edge weights were distance-dependent. The final layer is sum-pooled into an embedding. This embedding is then passed through two (independent) multilayer perceptrons to predict two outputs: the ANPR pose quality score, and the Vina docking score. Those outputs are then concatenated to the embedding and passed through a third multilayer perceptron which outputs the predicted pKi.

More details about the method and parameters can be found in (Gniewek et al., n.d.; Stafford et al., 2022).

### MSM-Weighting of Docking and pKi Predictions

To determine an overall MSM-weighted pKi prediction from pKi predictions for each MSM state, we first selected a single highest scoring pose for each state based on the AtomNet PoseRanker predictions. Next, we converted the predicted pKi value to an association constant. Then, we found a macro-association constant from the individual mico-association constants:

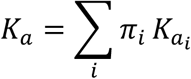

We use association constants because this ensures that large contributions to the sum come from states with either a high equilibrium probability, a large association constant (i.e., favor ligand binding), or both. States that have small association constants or low equilibrium probabilities will have a minimal contribution to the overall association constant. Finally, we convert the overall association constant to a pKi by taking the - log10 of its inverse.

For docking scores which are in units of kcal/mol, we follow a similar procedure. Given there were multiple poses for each MSM state, we selected the pose with the highest ANPR prediction for that state. Docking scores are then converted to association constants:

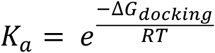

Then we follow the same aggregation procedure:

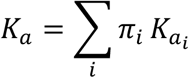

Finally, we convert this overall association constant into a pKi by taking the -log10 of its inverse.

## Acknowledgements

We would like to thank Atomwise for financing this project. AM was also supported by the National Institutes of Health F30 Fellowship (1F30HL162431-01A1). GRB holds a NSF grant MCB 2218156 (GRB) and NIH grants R01 GM124007 (GRB) and RF1AG067194 (GRB). GRB holds a Packard Fellowship for Science and Engineering from The David & Lucile Packard Foundation.

## Notes

AtomNet is a registered trademark of Atomwise Inc.

