## Supplement for "Discovery of a cryptic pocket in the AI-predicted structure of PPM1D phosphatase explains the binding site and potency of its allosteric inhibitors"

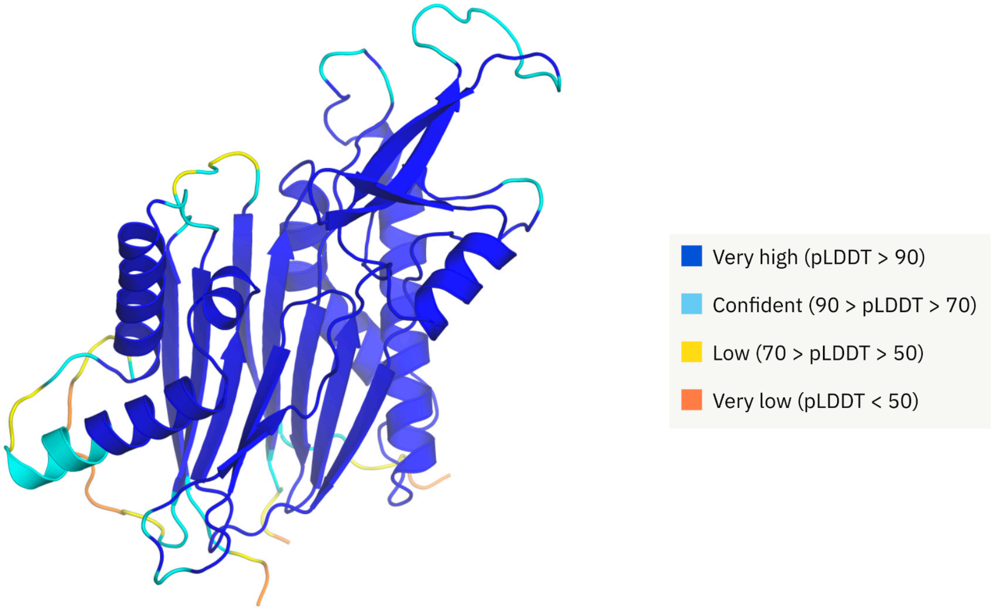


**Fig S1:** The AF predicted structure for Wip1 reveals high confidence prediction in the flap domain. Unlike other homology models, AF predicts that the flap domain contains ordered $\beta$-strands.

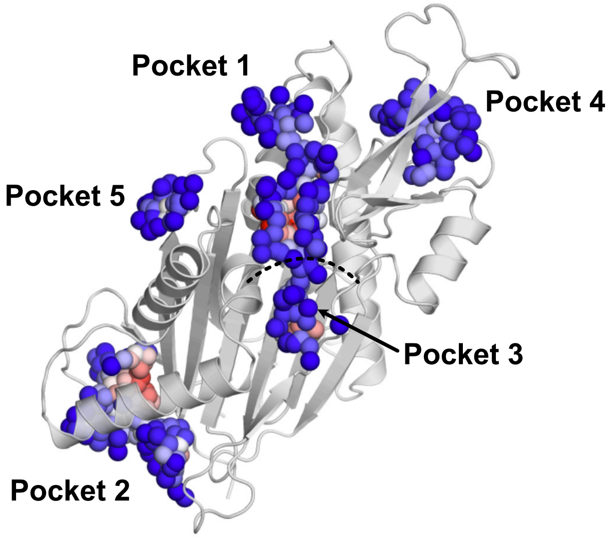


| name | rank | score | probability |
|---|---|---|---|
| pocket1 | 1 | 35.51 | 0.892 |
| pocket2 | 2 | 27.07 | 0.825 |
| pocket3 | 3 | 9.44 | 0.339 |
| pocket4 | 4 | 8.62 | 0.296 |
| pocket5 | 5 | 3.42 | 0.062 |

**Fig S2:** P2Rank predictions for the AlphaFold Wip1 structure reveal that there are no high scoring pockets at the flap or hinge. Spheres represent pocket grid points and are colored by their probability of being a ligand-binding site (red is higher, blue is lower). Only pocket 1 and 2 are predicted to bind ligands. Pocket 1 is found at the active site while pocket 2 has not been implicated as a potential binding site in previous work. The pocket involving the flap domain (pocket 4) is low scoring while there are no high-scoring pockets at the hinge.

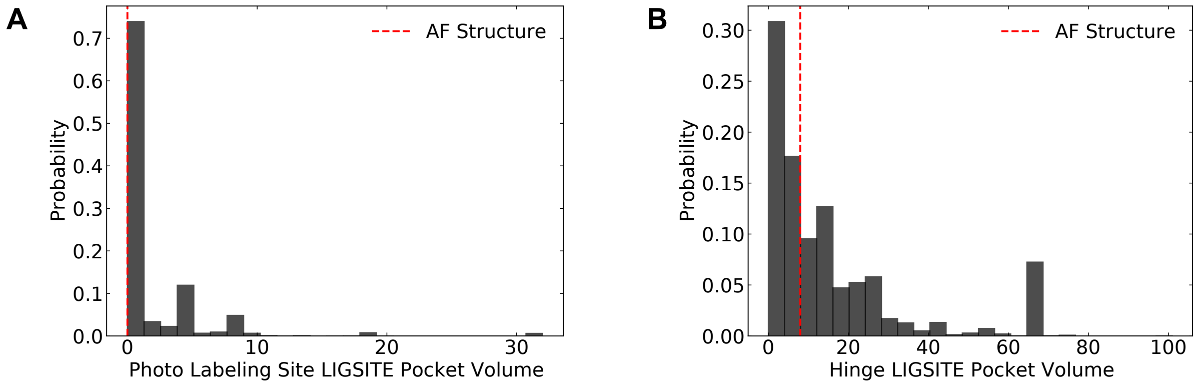


**Fig S3:** LIGSITE pocket volumes calculated across MSM at proposed binding sites show that cryptic pockets form at both the photolabeling site and hinge pocket volumes. At both these sites, the pocket volumes in the AF structure (red arrow) are much smaller than the maximum sampled in the simulations. We calculated LIGSITE pocket volumes for all structures (see Methods) and restricted pocket elements that were within 5 Angstroms of select residues. The photolabeling site was defined as residues 219 and 236 while the hinge residues were defined as residues 155 to 166.

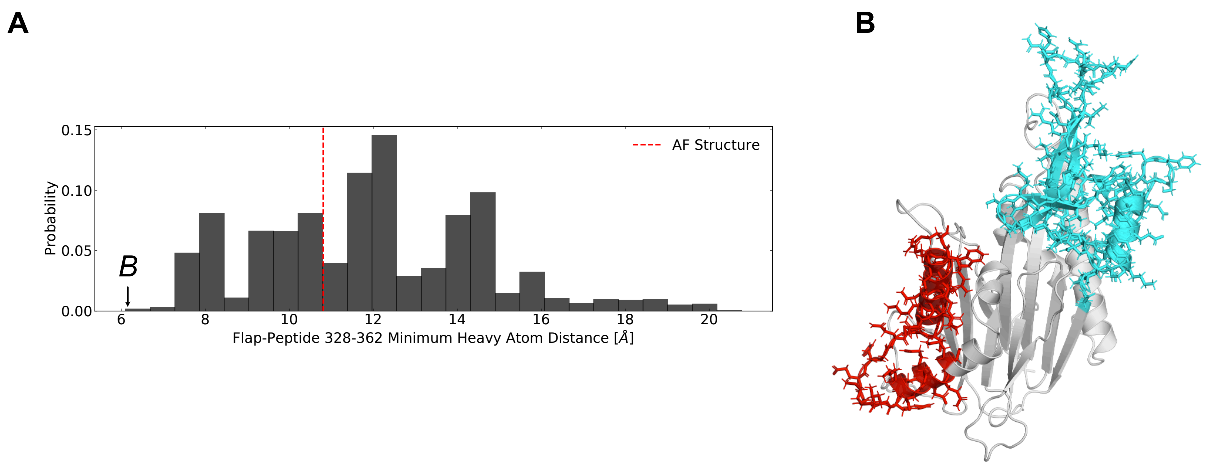


**Fig S4:** The flap domain approaches residues 328-362 in simulation despite being separated by over 10 Angstroms in the AlphaFold structure. A) Equilibrium probability-weighted distribution of flap domain minimum heavy atom distance to residues 328-362 shows that the flap domain can extend to this region. B) Wip1 structure with the smallest heavy atom distance between the flap and residues 328-362. Val241 from the flap is in proximity to Trp358 in this structure.

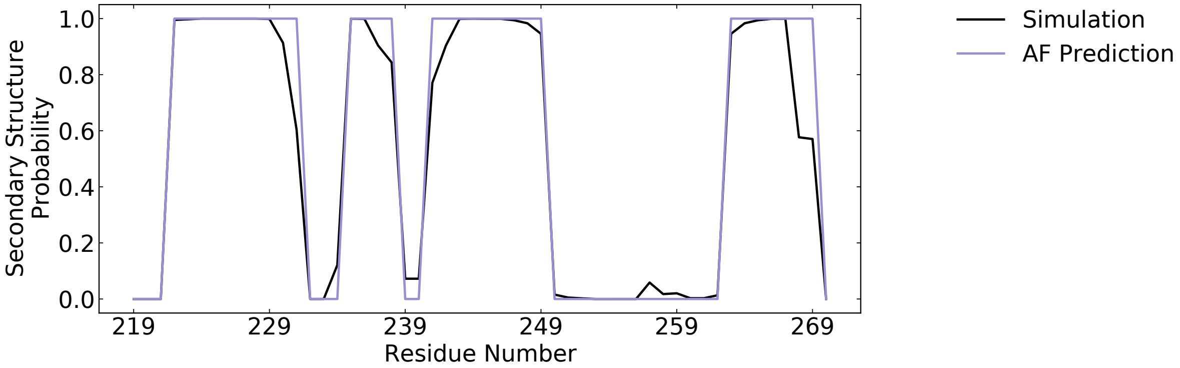


**Fig S5:** The β-strands in the flap domain remain stable throughout the simulations as predicted by AlphaFold. We calculated the equilibrium distribution of secondary structure elements for each residue using the dictionary of protein secondary structure (DSSP) assignments. Besides the end of the β-strand spanning residues 263-269, each β-strand has an equilibrium probability of being in a non-coil assignment near 1.0 (black line). The purple line shows the assignments for the AF-predicted starting structure.

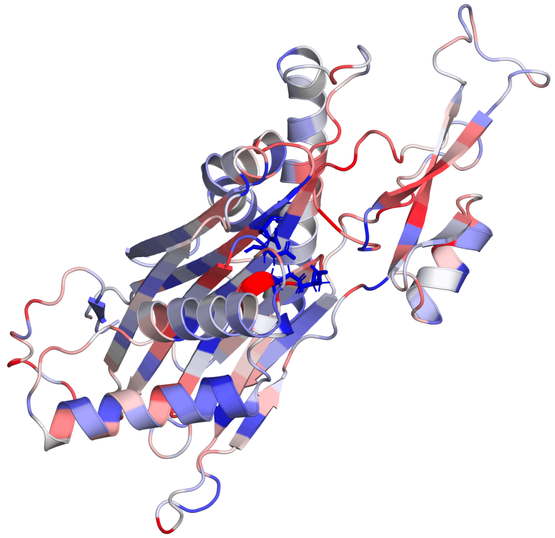

**Fig S6:** The Wip1 AlphaFold structure colored by the maximum residue-level increase in ligand-binding probability (red is higher, blue is lower) shows that the flap domain is a hotspot for increases in ligand-binding probability.

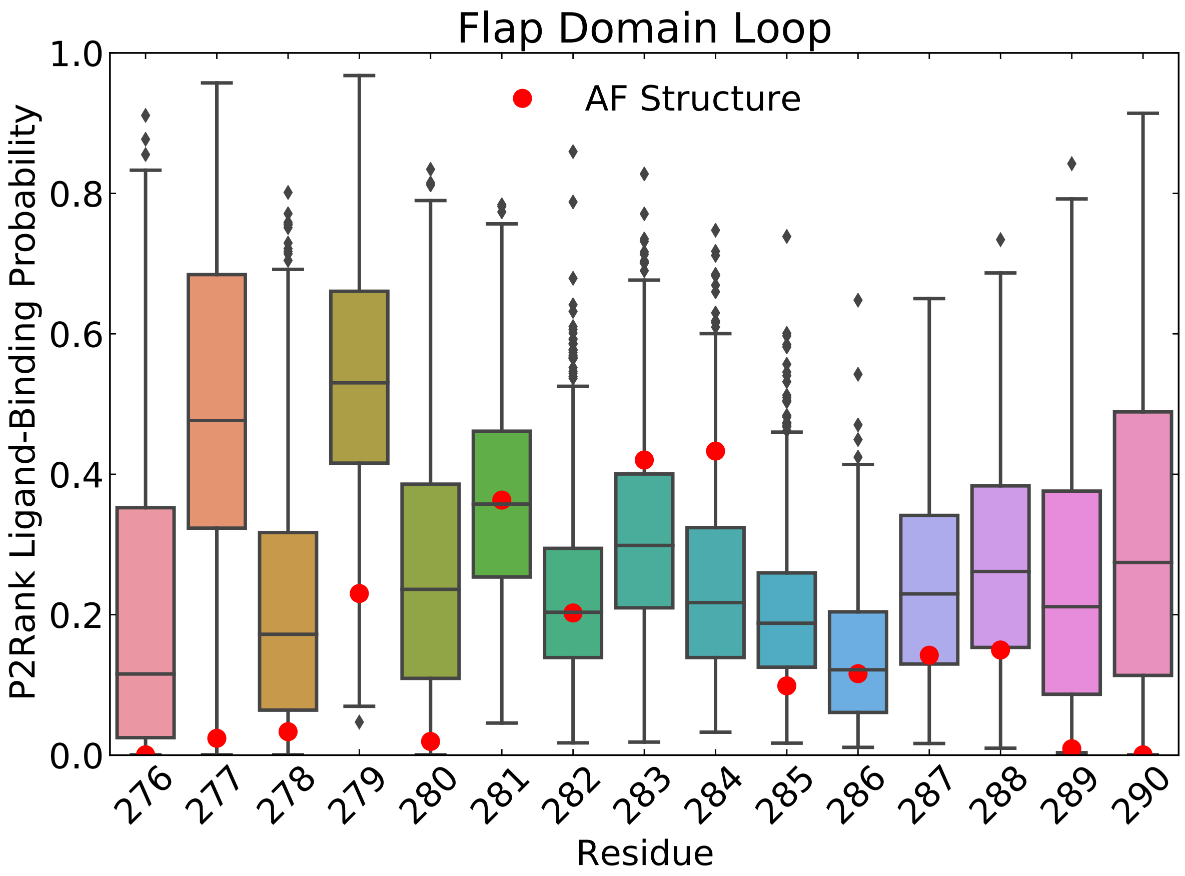


**Fig S7:** The flap domain loop spanning residues 276-290 is a hotspot for increases in the P2Rank ligand-binding probability. Box-and-whisker plots show the distribution of residue-level ligand-binding probabilities across the Wip1 MSM cluster center structures. The red dot indicates the predicted ligand-binding for the AF structure. Nearly all residues in this region exceed 0.6 at some point in the simulation.

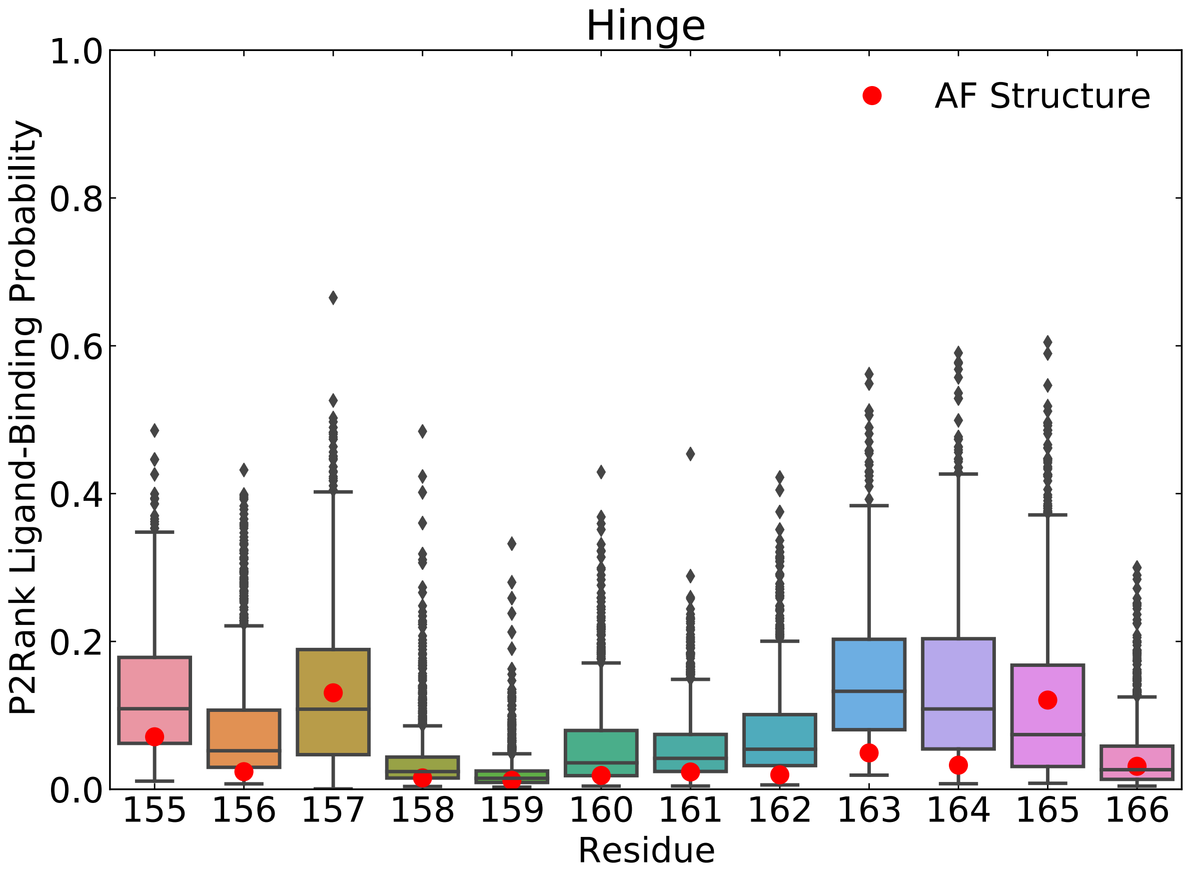


**Fig S8:** The hinge domain does not achieve high P2Rank ligand-binding probabilities at any point in the simulation. Box-and-whisker plots show the distribution of residue-level ligand-binding probabilities across the Wip1 MSM cluster center structures. The red dot indicates the predicted ligand-binding for the AF structure. Nearly all residues in this region do not exceed 0.6 at any point in the simulation.

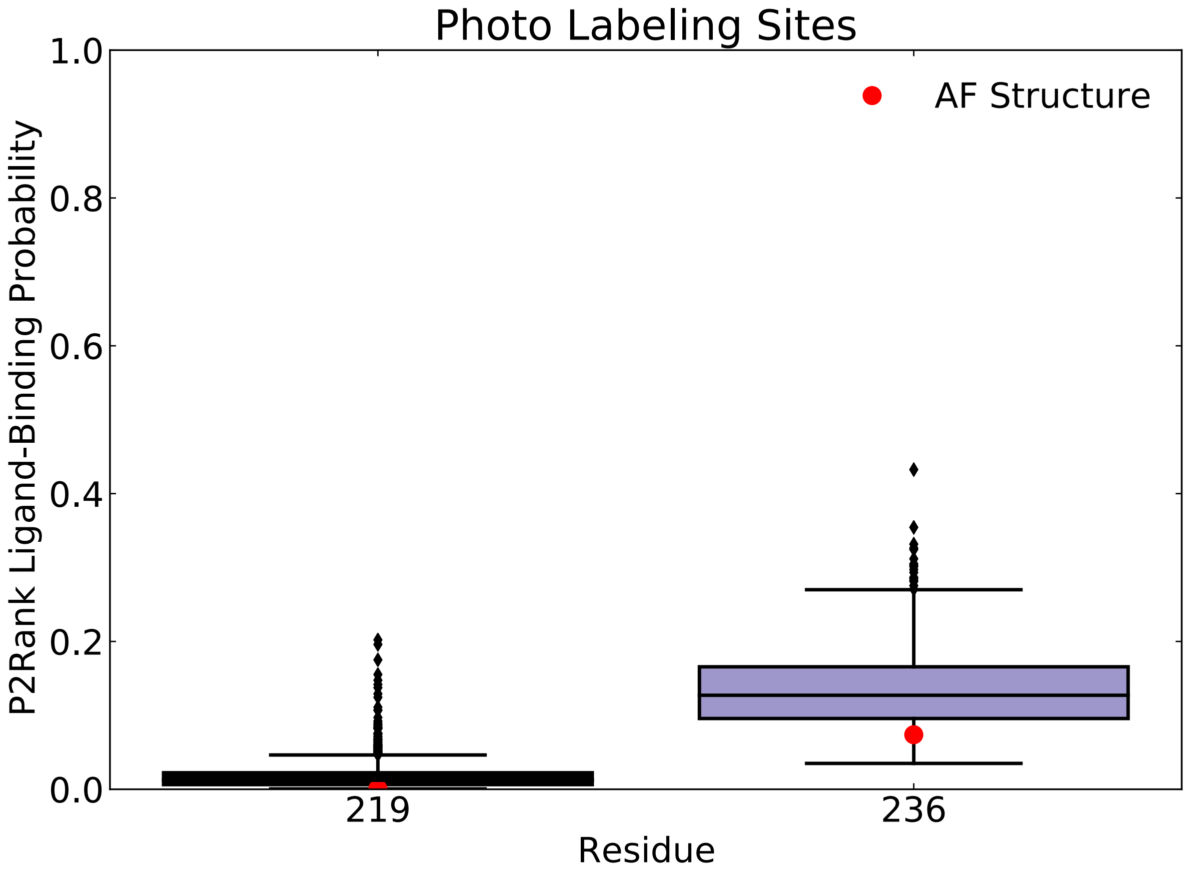


**Fig S9:** The photolabeling sites (P219 and M236) do not achieve high P2Rank ligand-binding probabilities at any point in the simulation. Box-and-whisker plots show the distribution of residue-level ligand-binding probabilities across the Wip1 MSM cluster center structures. The red dot indicates the predicted ligand-binding for the AF structure.

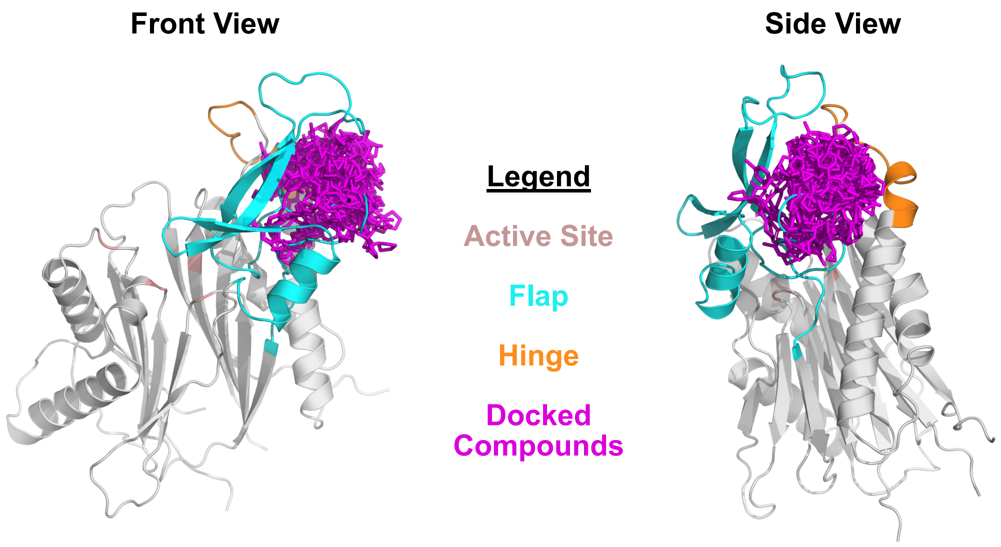


**Fig S10:** An overlay of docked poses that receive poseNet predictions greater than 0.5 shows that high quality poses cluster between the flap and hinge. The AlphaFold Wip1 structure is depicted, and poses may clash with this structure since these compounds were docked to other structures in the Markov State Model.

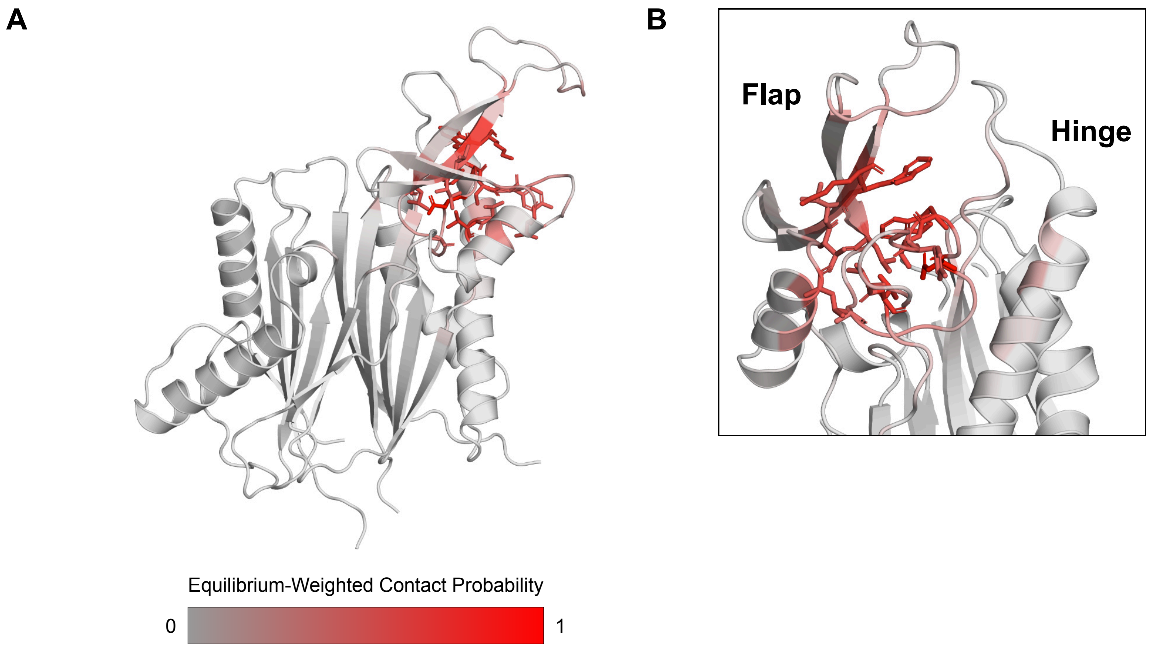


**Fig S11:** Equilibrium-weighted contact probabilities implicate the flap as the main site of GSK compound binding. Given multiple poses for each structure in the Wip1 MSM, we selected the pose with the highest poseNet prediction as the most likely binding mode for that state. We then designated any residues within 4 Angstroms of a ligand heavy atom as ones that made with the docked ligand. B) Inset showing a zoomed-in side view of the flap and hinge. Residues D277, W279, and V290 were most likely to contact the best pose in each state.

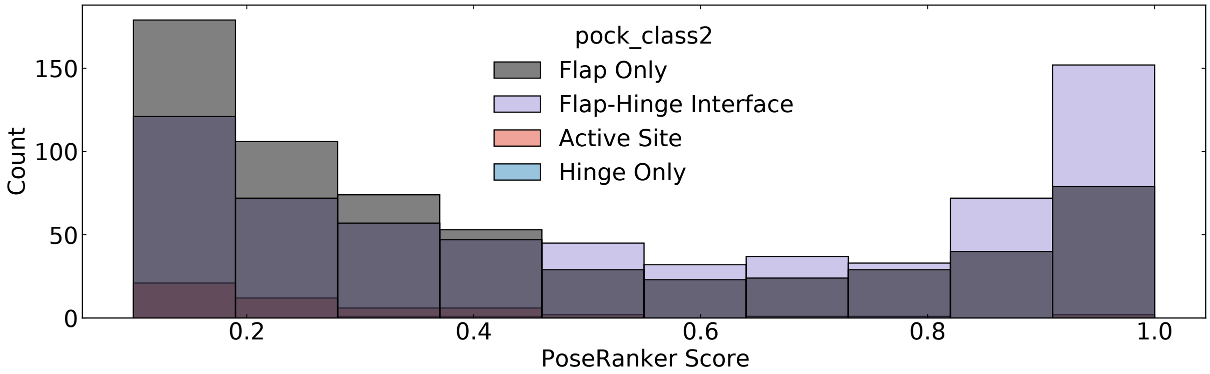


**Fig S12:** The distributions of the AtomNet® PoseRanker scores per binding site shows that poses at the flap-hinge interface receive higher scores. Scores below 0.1 are not shown to better visualize higher scores.

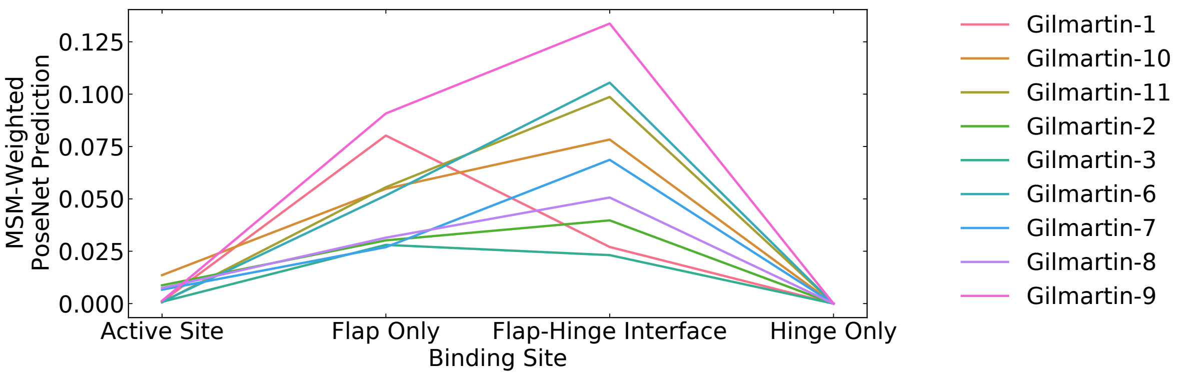


**Fig S13:** MSM-weighted PoseRanker predictions per binding site and compound shows that the flap-hinge interface is the highest-scoring binding stie for all but two compounds. The lowest-scoring compound is compound 3, which has no activity.

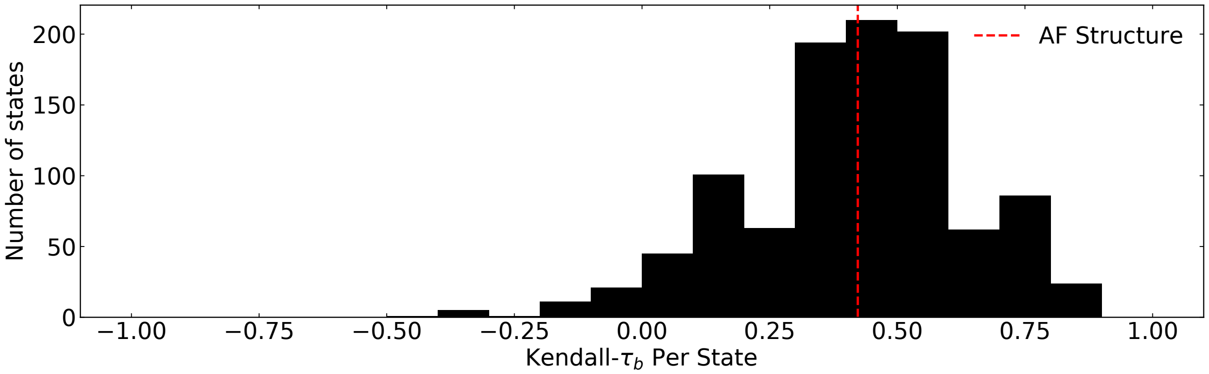


**Fig S14:** Histogram of tau-b scores for each MSM cluster center show that the ability to rank compounds is sensitive to the structure used for docking.

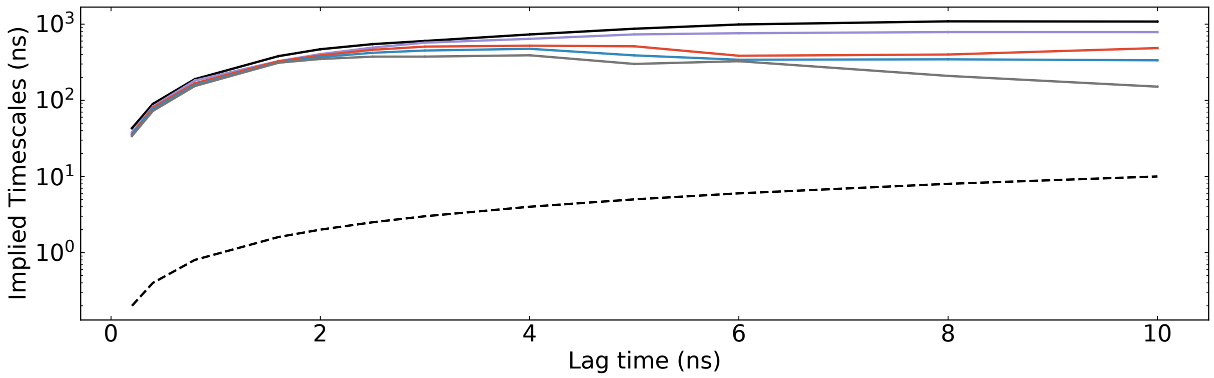


**Fig S14:** Implied timescales plot shows convergence on logarithmic scale around 2 ns.

**Table S1:** The AtomNet® PoseRanker AlphaFold Structure Predictions

| **Compound** | **PoseRanker Highest Score** |
|---|---|
| Gilmartin-1 | 1.28E-04 |
| Gilmartin-10 | 8.82E-07 |
| Gilmartin-11 | 4.86E-05 |
| Gilmartin-2 | 3.83E-05 |
| Gilmartin-3 | 2.29E-06 |
| Gilmartin-6 | 3.81E-05 |
| Gilmartin-7 | 4.33E-04 |
| Gilmartin-8 | 8.67E-02 |
| Gilmartin-9 | 2.21E-04 |
